# In computer explore The neutralization mechanism of Amubarvimab and Romlusevimab against SARS-COV-2 mutants

**DOI:** 10.1101/2024.09.14.613008

**Authors:** Xinkang Huan, Hongwei Gao

**Affiliations:** School of Life Science, Ludong University, Yantai, Shandong, 264025, China

**Keywords:** COVID-19, SARS-CoV-2, mutations, RBD, Amubarvimab, Romlusevimab, computer method

## Abstract

Since the end of 2019, The coronavirus disease 2019 (COVID-19) has been endemic worldwide for three years, causing more than 6.95 million deaths and having a massive impact on the global political economy. With time, the Severe acute respiratory syndrome coronavirus 2(SARS-COV-2) is also constantly mutating. Mutations lead to stronger infectivity or virulence of the virus, and some monoclonal antibodies against wild-type SARS-COV-2 are also challenging to play a role. Amubarvimab and Romlusevimab were originally developed against wild-type SARS-COV-2; however, these monoclonal antibodies’ neutralizing efficacy and mechanism against these mutants are unknown. In this study, the binding ability of Amubarvimab and Romlusevimab to 7 mutant strains were tested by computer method and the interaction mechanism was explored. Our experimental data show that Amubarvimab can effectively bind most mutations and maintain the stability of the complexes mainly through hydrogen bond interaction; However, the binding efficiency of Romlusevimab was lower than that of Amubarvimab, and the stability of the complexes was maintained mainly through electrostatic interaction. Both Amubarvimab and Romlusevimab show low binding potency against E406W and Q498Y mutations, so there is a certain probability of immune escape in the face of variants carrying E406W and Q498Y mutations when Amubarvimab and Romlusevimab are used in combination.

## 1. Introduction

At the end of 2019, SARS-COV-2 was first detected in Wuhan, China(Hu et al., 2020). Three years later, traces of SARS-COV-2 are still spreading worldwide. The COVID-19 caused by the SARS-COV-2 infection has caused an enormous disaster to the world (6.95 million people have died as of September 2023) and seriously threatened the stability of the global political economy(Duma et al., 2022). Although countries worldwide have slowed down their policies on the SARS-COV-2 epidemic, the repeated outbreaks caused by the SARS-COV-2 variants are still causing great distress to people(Salzberger et al., 2020).

The SARS-COV-2 is a betacoronavirus, positive-sense single-stranded RNA virus with a gene length of about 30kb(Yang and Rao, 2021;Si et al., 2023). From the 5 ‘end to the 3’ end, two polymeric protein precursors pp1a pp1ab, spike protein (S), membrane protein (M), envelope protein (E), and nucleocapsid protein (N) are encoded successively. The SARS-COV-2 is spherical, about 120nm, consisting of a coat protein and RNA, with a phospholipid bilayer and spike protein on the surface(Hasöksüz et al., 2020;Hu et al., 2020;Yadav et al., 2021;Ravi et al., 2022). Spike protein is the main element of virus invasion cells; it is a homologous trimer formed by non-covalent bonding of three identical subunits. Each subunit of S protein is composed of 1273 amino acids(Schaub et al., 2021). The host protease cuts the S protein into S1 and S2 subunits when the virus infects cells. The S1 subunit is responsible for the specific recognition of the angiotensin-converting enzyme 2 (ACE2) receptor; The S2 subunit is responsible for fusing with the host cell membrane. The S1 subunit contains the N-terminal domain (NTD) and the C-terminal receptor (CTD)(Gusev et al., 2022). SARS-COV-2 interacts with hACE2 through the CTD domain of S1. The conformation of the S1 subunit will change, resulting in binding with ACE2 and then shedding with ACE2; at this point, the S2 subunit becomes a stable fusion state with the host cell membrane. The S2 subunit contains a fusion peptide (FP), heptapeptide repeats (HR1 and HR2), transmembrane domain (TM), and cytoplasmic domain (CP). The S2 subunit changes its conformation by inserting fusion peptides into the host cell. HR1 and HR2 spiral compress to form a tight six-helix bundle (6HB), resulting in the fusion of the viral membrane with the cell membrane. The nucleic acid molecule is released into the cell to complete the infection(Jackson et al., 2022;Marcink et al., 2022;Mehra and Kepp, 2022). The receptor binding domain (RBD) is a critical component of the S protein subunit (S1) that binds to angiotensin-converting enzyme 2 (ACE2), a recognized viral entry receptor. Because RBD-ACE2 binding occurs at the origin of infection, RBD is a prime target for developing therapeutic interventions(Muhammed et al., 2021). Therapeutic mechanisms aimed at neutralizing viral infection include inhibiting the interaction between the spike proteins RBD and ACE2 or disrupting the S2 domain activity of the S protein to prevent membrane fusion(Barnes et al., 2020;Hwang et al., 2022).

Due to the instability of single-stranded RNA, many variant strains of SARS-COV-2 have emerged worldwide, and these mutations have enabled the SARS-COV-2 to gain evolutionary advantages, some of which occur in the RBD region of spike proteins, resulting in the SARS-COV-2 variants with high virulence and high infectivity(Sanjuán and Domingo-Calap, 2016;Flores-Vega et al., 2022). E406W, N439K, L452R, Y453F, F486L, Q498Y and N501 occurred in the region of the RBD mutations; some of these mutations showed high affinity for the ACE2 receptors more quickly than the parent virus amplification; some showed higher virulence than their parent viruses; some are beneficial to maintain the conformation of the virus infection state and improve the efficiency of the virus entering target cells; others hide the neutralizing epitopes of monoclonal antibodies to enable immune escape reactions(Motozono et al., 2021;Sanyaolu et al., 2021;Thomson et al., 2021;Addetia et al., 2022;Erausquin et al., 2022;Pondé, 2022).

Amubarvimab and Romlusevimab, a combination of antibodies developed by Brii Biosciences, Tsinghua University and the Third People’s Hospital of Shenzhen, were first introduced in China. To treat high-risk or dying COVID-19 patients over 18 years or 12-17 years weighing more than 40kg. (Hoy, 2022;Liu et al., 2022). Amubarvimab and Romlusevimab are human IgG1 monoclonal antibodies isolated from B cells of patients with COVID-19. They can specifically against the wild-type SARS-CoV-2 and do not neutralize other coronaviruses. Amubarvimab and Romlusevimab target different epitope regions in RBD in coronavirus spike glycoproteins in a non-competitive manner, allowing the combination to maintain neutralizing activity against several SARS-COV-2 variants(Ju et al., 2020;Ji et al., 2022;Liao et al., 2022). Compared with monoclonal antibodies composed of a single antibody, mixed antibodies have a higher barrier to prevent immune escape from the SARS-COV-2, as at least two mutations of the RBD amino acid are required(Hurt and Wheatley, 2021).

In this experiment, we conducted molecular docking of Amubarvimab and Romlusevimab with several RBD mutants mentioned above, aiming to screen the mutations that can be effectively neutralized by monoclonal antibodies and explain the neutralization mechanism of monoclonal antibodies.

## 2. Experimental section

### 2.1 Data Mining

Amubarvimab and Romlusevimab were searched from the PROTEIN DATA BANK RCSB PDB: Homepage database to find the structure of the two antibodies and the wild-type SARS-COV-2 protein (8GX9). The mutation information of SARS-COV-2 RBD was obtained in NCBI Virus. The SAES-COV-2 RBD mutant strains include the following seven types: E406W, N439K, L452R, Y453F, F486L, Q498Y, N501Y.

### 2.2 Build And Edit RBD

Use the build and Edit Protein program in Discovery Studio and mutation of wild-type RBD: E406W, N439K, L452R, Y453F, F486L, Q498Y, N501Y. The Build and Edit Protein tools allow you to build and modify peptides and proteins quickly. These tasks are supported mutating residues, mutating one or more selected residues to a specific residue type.

### 2.3 Molecular Docking

Computer simulation of molecular docking technology can screen potential small molecule ligand; active drug molecules; drug action targets and simulate protein interaction patterns, which is helpful to accelerate the process of drug development(Agrawal et al., 2019). In the study of protein binding, the complex model, which is difficult to resolve by traditional methods(Co-immunoprecipitation, Enzyme-linked immunosorbent assay), can be used to dock the protein’s three-dimensional structure by molecular docking technology to predict the binding mode of the two. In order to better elucidate the biological and molecular mechanism of Amubarvimab and Romlusevimab in neutralizing the SARS-COV-2 mutant strains, this paper adopts Discovery Studio 2020 software to conduct molecular docking simulation calculations on the SARS-COV-2 receptor binding domain RBD of Amubarvimab and Romlusevimab(Pierce et al., 2014). According to docking scores and conformation analysis, the optimal binding mode of the complex was screened to promote the understanding of the structure and function of the SARS-COV-2 RBD, develop corresponding neutralizing antibodies of the SARS-COV-2, and understand the neutralization of Amubarvimab and Romlusevimab against the SARS-COV-2 strains. ZDOCK program was used for molecular docking. ZDOCK is a rigid protein docking technology based on Fast Fourier Transform, which can cluster conformations according to ligand positions and order conformations according to ZRANK scores(Chen et al., 2003).

### 2.4 Molecular dynamics simulation

Molecular dynamics simulations of Amubarvimab – wild type novel coronavirus complex and Romlusevimab – wild type novel coronavirus complex were performed to verify the conformation stability of the complex. Molecular dynamics simulation, referred to as MD simulation, was developed in 1966, and with the development of computer technology and rapid development, MD is now mostly used in the study of protein stability, protein folding and superposition interaction, protein and ligand interaction, and molecular recognition and other fields. We used AMBER18 to perform A molecular dynamics simulation in which the RD-MAB complex structure obtained by molecular docking was assigned a force field to the protein ff14SB and placed in a water box with a truncated octahedron at a buffer distance of 10 A. Na+ or Cl-ions are added to the water box to neutralize the net charge of the system, keeping the net charge of the system at 0. Then, energy minimization, system heating, system equilibrium and molecular dynamics simulation were carried out successively. The energy minimization stage was completed by the 10,000-step conjugate gradient method and 10,000-step steepest descent method successively. After the energy minimization was completed, the system temperature was gradually increased to 310K within 300ps at a constant volume. Then the equilibrium simulation of 500ps at constant temperature and constant pressure was performed at 310K, and the molecular dynamics simulation of the whole complex system was performed at 200ns after rebalancing.

### 2.5 Epitope Analysis

Antigen epitopes are small sites on RBD that monoclonal antibodies can specifically bind(Burkovitz and Ofran, 2016;Wong and Jin, 2017). Amino acid residues at the mAb binding site interact with amino acid residues on RBD epitopes to form non-covalent bonds and provide necessary energy for the stability of the complex(Guerois et al., 2002). Visualization of epitopes plays an essential role in elucidating the interaction mechanism and molecular recognition mechanism between antigens and antibodies(Nelson et al., 2000). In this experiment, Pymol software was used, combined with the results of DS molecular docking, to label the primary structure and tertiary structure of RBD of each SARS-COV-2 variant by epitope amino acids, so as to compare the changes of RBD binding epitopes among different SARS-COV-2 variants.

### 2.6 Analyses Protein-Protein Interactions

The optimal conformation of the complex was obtained by molecular docking technique, and then the conformation of the complex was analyzed. The quantitative calculation results of the hydrogen bond, electrostatic interaction and hydrophobic interaction between the mAbs and RBD complexes were obtained using the analysis protein interface program(Soleymani et al., 2022). The Analyze Protein Interface protocol provides methods for identifying the residues found at a protein-protein interface and analyzing the interactions that occur there. The interface residues can be identified as those within a specified cutoff distance of the neighboring chain or as the residues whose solvent accessible surface area is different depending on whether the protein chains are in a complex or are isolated. The establishment of this method shows that it is feasible to simulate protein interaction by computer, which provides an adequate theoretical basis for protein drug development, molecular recognition and antibody construction, and has a wide range of medical applications(Du et al., 2016;Lee et al., 2019).

## 3 Results and discussion

### 3.1 Amubarvimab and Romlusevimab can effectively neutralize wild-type SARS-CoV-2

In order to verify the feasibility of the experimental method, we first studied the binding mode of Amubarvimab and Romlusevimab with wild-type RBD. After comparison, we found that the complex model screened by this method was consistent with the results obtained by the previous experimenter through the X-ray diffraction method(Ji et al., 2022). This complex model was not only used to verify the feasibility of the test method but also served as a control group to compare the differences in the binding patterns between RBD and monoclonal antibodies of various SARS-COV-2 mutants. Through molecular docking technology, we explored the complexes structure of Amubarvimab and Romlusevimab with SARS-COV-2 RBD and found that they can effectively bind to the RBD of SARS-COV-2. In addition, Amubarvimab and Romlusevimab are bound to different epitopes of SARS-COV-2 RBD and effectively neutralize SARS-COV-2 through non-competitive binding. The binding regions of Amubarvimab mainly focus on 417-420,473-493, and the binding sites of ACE2 receptors have extensive overlap, indicating that Amubarvimab prevents SARS-COV-2 from invading cells through competitive binding. In contrast, Romlusevimab binds to the opposite side of RBD, with binding sites mainly concentrated in 340-360 and 440-460. The figure 1 shows the schematic diagram of Amubarvimab and Romlusevimab combined with RBD. The blue surface model is wild-type RBD, the orange cartoon model is Amubarvimab, and the green cartoon model is Romlusevimab.

**Figure 1.**
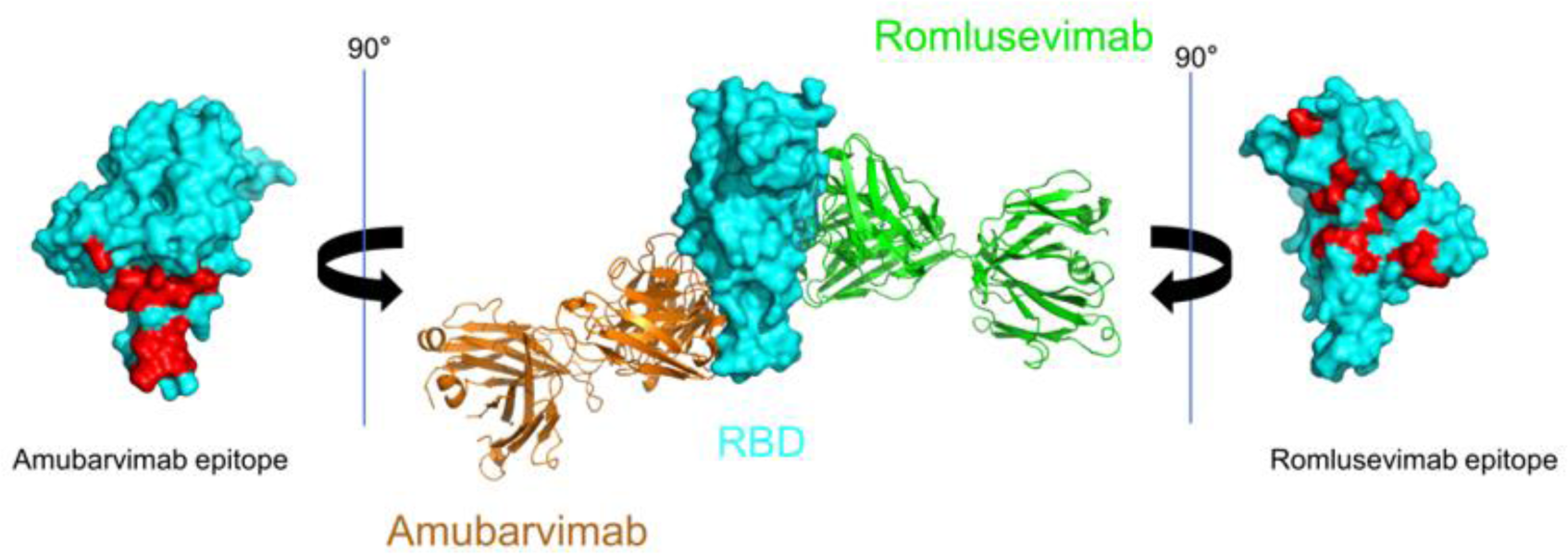
Schematic diagram of Amubarvimab-RBD-Romlusevimab binding. The blue surface model is the RBD of wild-type SARS-COV-2, the orange cartoon model is Amubarvimab, and the green cartoon model is Romlusevimab. When the middle wild-type SARS-COV-2 RBD was rotated 90° to the right, the neutralizing epitope of Amubarvimab on the left was obtained, and the neutralizing epitope of Romlusevimab on the right was obtained by rotating 90° to the left. The red areas are epitopes neutralized by monoclonal antibodies.

In the molecular dynamics simulation of 200ns, the overall stability of AMU-W complex was better. As shown in Figure 2, after 60ns, the entire complex system tended to equilibrium, and the overall RMSD value fluctuated around 1A. The stability of the ROM-W composite system was slightly lower than that of the AMU-W composite system. After 75ns, the ROM-W composite system tended to be stable, and the RMSD value fluctuated within 1.5 A, which was within the acceptable range. This indicates that both Amubarvimab and Romlusevimab can stably bind to wild-type novel coronavirus RBD, and the secondary and tertiary structures of the complex are also very stable.

**Figure. 2.**
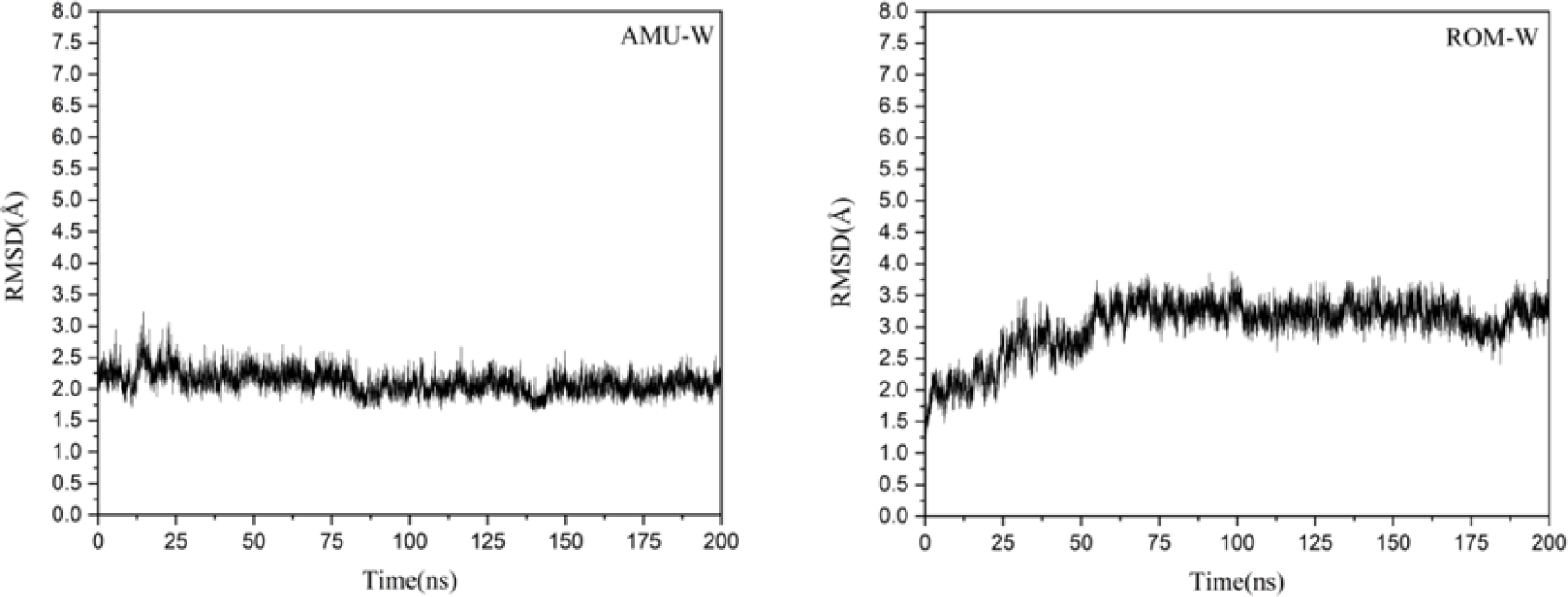
RMSD of the mAb-RBD complex

As shown in Table 1, the binding free energies of Amubarvimab and Romlusevimab with wild-type RBD were −48.35kcal mol-1 and −55.99kcal mol-1, respectively, and the energy of AMU-W complex system was slightly higher than ROM-W, indicating that the neutralizing effect of Romlusevimab was stronger than that of Amubarvimab. By analyzing the contribution value of each energy to the total binding energy, it is found that in the complex ROM-W system, the electrostatic interaction is −475.07kal mol-1, which accounts for the largest proportion and plays the main binding capacity. It is speculated that there are many charged amino acid residues at the binding interface, which contribute energy to the binding of the complex system.

**Table 1.**
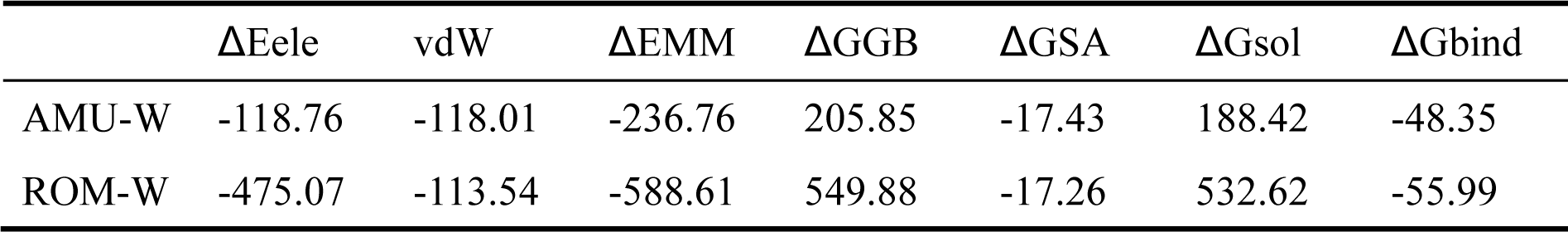
Binding energy of Amubarvimab and Romlusevimab antibodies to wild-type RBD.

### 3.2 Antibody binding to other mutant RBD was determined

Since the single point mutation had little effect on the overall conformation of the complex, molecular dynamics simulations of other complexes were not performed. We then tested all possible docking results between 2 monoclonal antibodies and all mutated RBD in a computer. A total of 14 RBD-mAb protein complexes were obtained using Discovery Studio. The Wild-mAb was used as the control group, and docking scores were ranked. Table 2 shows the docking scores between Amubarvimab and various RBDs, among which the Wild-Amubarvimab complex has the highest docking score and the E406W-Romlusevimab complex has the lowest docking score. We found that Amubarvimab had a high docking score with most RBD variants, among which the docking score with wild-type RBD was the highest, and only E406W and Q498Y mutations showed relatively low docking scores. So we hypothesized that Amubarvimab could effectively neutralize wild-type SARS-COV-2, can effectively neutralize the L452R, N439K, Y453F, F486L, N501Y mutation of SARS-COV-2 variants, and effectiveness of E406W, Q498Ymutation; On the contrary, the docking score of Romlusevimab and various RBD variants was relatively low, which was also in line with previous experimental verification, among which the docking score of E406W was the lowest, followed by Q498Y. We can guess Romlusevimab neutralization contains N439K, L452R, Y453F, F486L and N501Y mutation of SARS-COV-2 variants, but lower than neutralizing efficacy Amubarvimab, cannot effectively neutralize E406W, Q498Y both variants. Docking score is a preliminary assessment of the neutralization efficacy of monoclonal antibodies against SARS-COV-2 variants. To verify our hypothesis and obtain the molecular mechanism of RBD-mAb protein complexes interaction, we further improved our experiment: With wild-type RBD-MAB complexes as the control group, two complexes with high and two complexes with low bond scores were screened from RBD-Amubarvimab and RBD-Romlusevimab groups, respectively, and protein interaction analysis was performed on a total of 10 protein complexes.

**Table 2.** Docking score between monoclonal antibody and RBD.

| RBD with<br>mutation | Amubarvimab | Romlusevimab |
| --- | --- | --- |
| Wild RBD | 17.42 | 15.08 |
| E406W | 14.00 | 11.34 |
| N439K | 15.74 | 15.92 |
| L452R | 16.06 | 14.58 |
| Y453F | 15.02 | 15.08 |
| F486L | 14.92 | 14.50 |
| Q498Y | 13.44 | 11.70 |
| N501Y | 15.68 | 14.28 |

### 3.3 Epitope Analysis Results

We also analyzed neutralizing epitopes of RBD. Figure 3 shows wild-type and various mutant RBD epitopes when neutralized by Amubarvimab. It can be seen from the figure that Amubarvimab binding epitopes are concentrated, mainly in the bottom half of RBD (red, yellow and orange are neutralizing epitopes). The protein primary structure comparison analysis gives us a deeper understanding of binding epitopes. Table 3 shows the amino acid sequence of wild type and various mutant RBD interacting with Amubarvimab. From the table, we can see that Mainly concentrated in 3 regions 415-421, 450-460, and 470-490. Figure 4 shows the epitopes of wild type and various mutant RBD when Romlusevimab neutralizes them. Unlike Amubarvimab, epitopes in Romlusevimab are relatively dispersed and discontinuous, indicating that it is challenging to have centralized binding non-covalent forces at the molecular interaction level, which may also be the reason for the weak neutralizing efficacy of Romlusevimab. Subsequently, we also conducted the protein primary structure comparison analysis of various RBD combined with Romlusevimab. Table 4 shows the amino acid sequences that interact with mAb when the wild type and various mutant RBD are neutralized by Romlusevimab, which are mainly concentrated in two regions: 340-361, 448-470. After protein primary structure comparison, we found that the epitopes of Amubarvimab and Romlusevimab do not conflict, which is also consistent with the fact that Amubarvimab and Romlusevimab are non-competitive.

**Figure. 3.**
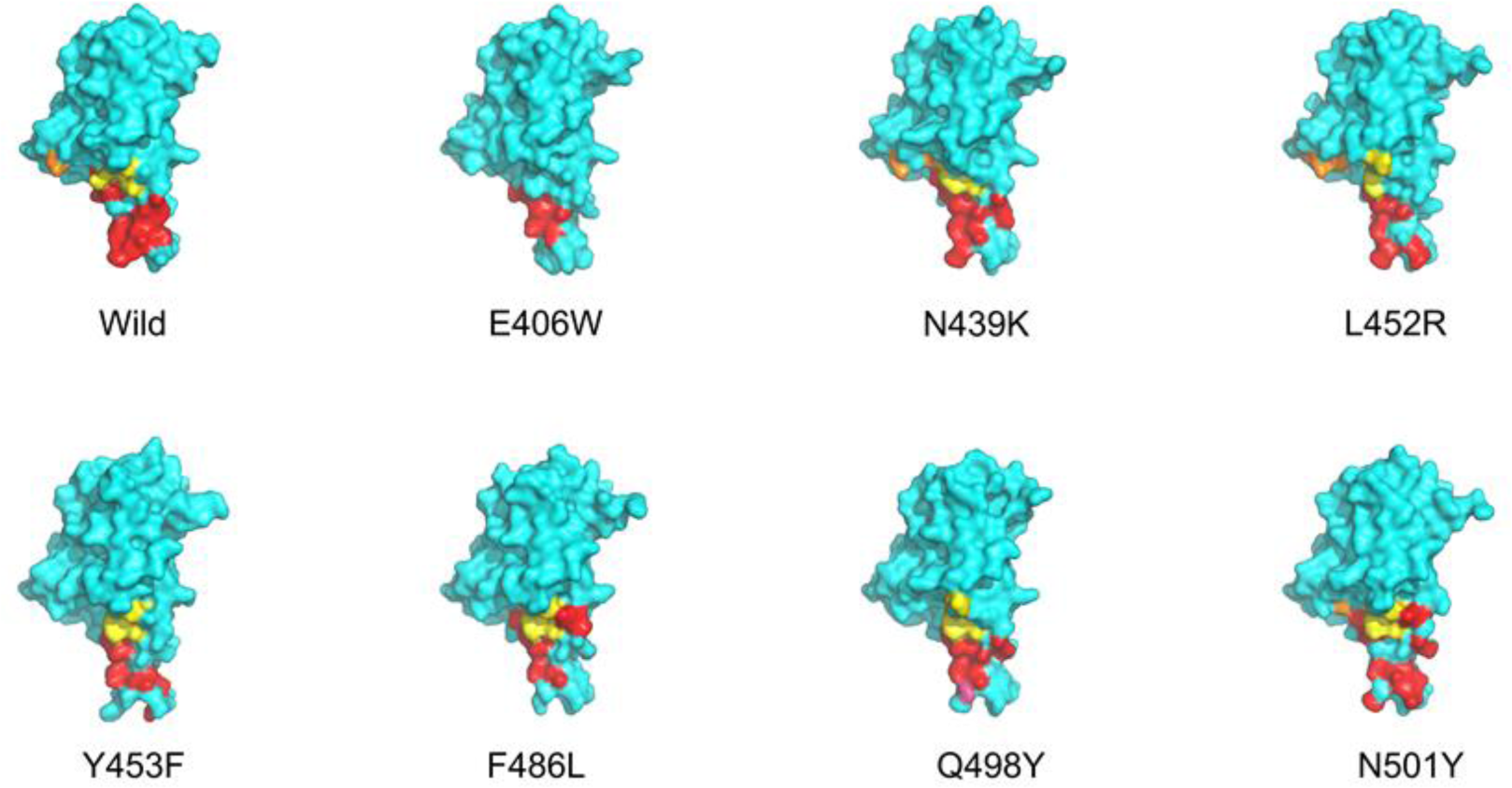
Epitope of wild-type SARS-COV-2 RBD and various mutant SARS-COV-2 RBD when neutralized by Amubarvimab. The blue surface models are various SARS-COV-2 RBD. Epitope 415-421 regions are shown in yellow, epitope 452-460 and epitope 471-493 areas are shown in red, and sites with low frequency and unconcentrated distribution are shown in orange.

**Table 3.**
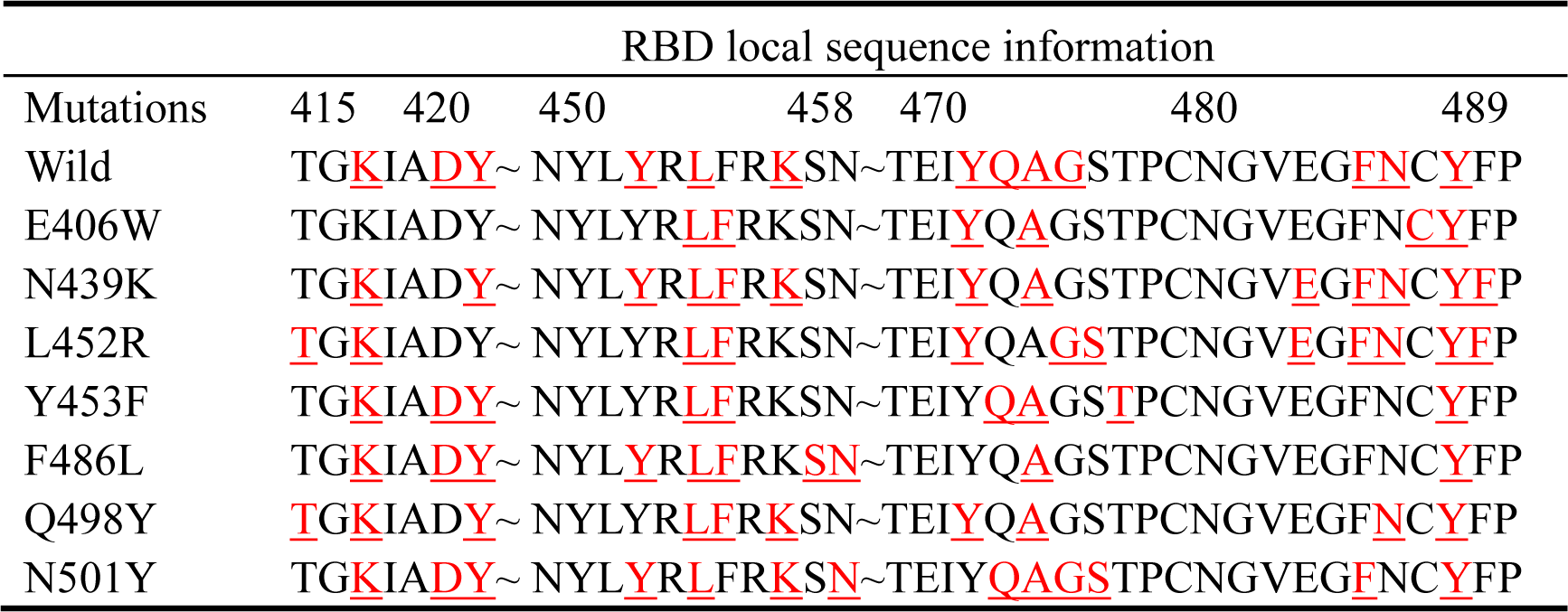
Local protein sequence information of RBD, epitopes that interact with Amubarvimab are highlighted in red.

**Figure 4.**
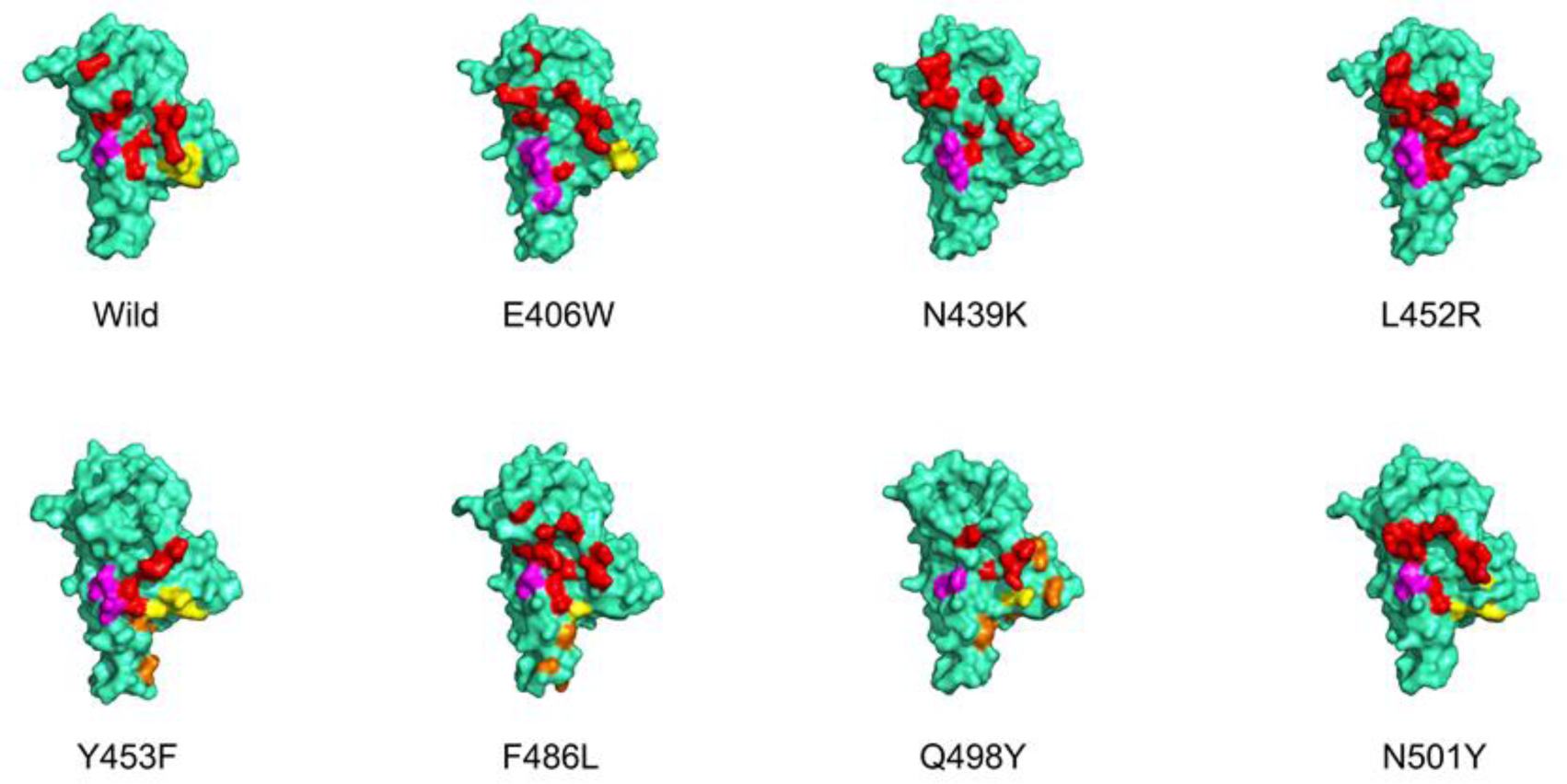
Epitope of wild-type SARS-COV-2 RBD and various mutant SARS-COV-2 RBD when neutralized by Romlusevimab. The green surface models are various SARS-COV-2 RBD. Epitope 340-361 region is shown in red, epitope 446-451 region is shown in yellow, epitope 466-470 area is shown in magenta, low frequency, distribution of the surface is shown in orange.

**Table 4.**
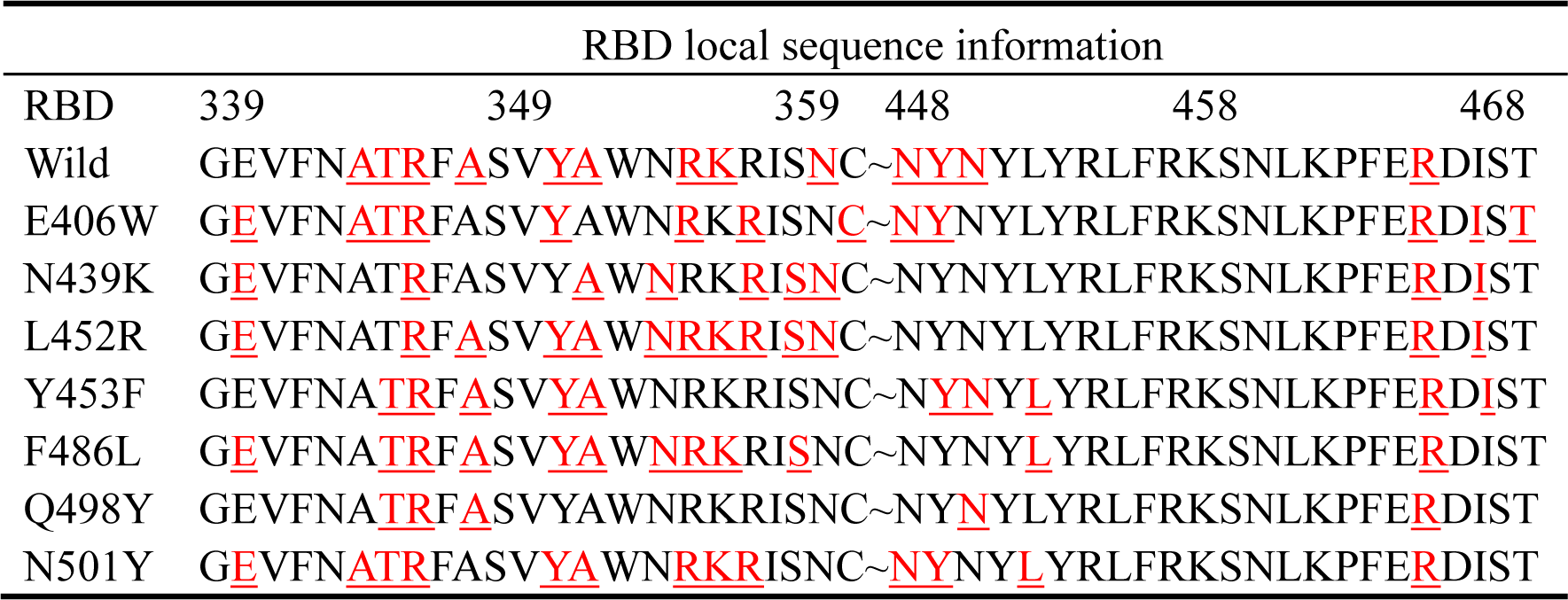
Local protein sequence information of RBD, epitopes that interact with Romlusevimab are highlighted in red.

### 3.4 Analysis of interaction between monoclonal antibody and RBD

By measuring the binding scores of monoclonal antibodies and each RBD, we selected five compounds from each group of RBD-mAb complexes, among which Wild RBD-Amubarvimab and Wild RBD-Romlusevimab were the control group. The binding scores were 17.42 and 15.08, respectively. N439K-Amubarvimab, L452R-Amubarvimab, N439K-Romlusevimab and Y453F-Romlusevimab complexes strongly bind protein complexes. The docking scores were 15.74, 16.06, 15.92 and 15.08 respectively. E406W-Amubarvimab, Q498Y-Amubarvimab, E406W-Romlusevimab and Q498Y-Romlusevimab complexes were weakly bound protein complexes, and their interworking scores were 14.00, 13.44, 11.34 and 11.70, respectively. According to docking scores, we found that the E406W mutation and Q498Y mutation showed lower docking scores for both Amubarvimab and Romlusevimab. Therefore, we speculated that mutant strains carrying E406W and Q498Y mutations might escape the immunosuppressive effects of Amubarvimab and Romlusevimab, or the neutralizing efficacy of the two monoclonal antibodies against E406W and Q498Y mutations would be significantly reduced.

Table 5 summarizes the participation of various non-covalent bonds among the complexes and the changes in covalent bonds. The Wild-Amubarvimab complex is the most stable, and it is stabilized by 16 Hydrogen bonds, 2 Electrostatic and 6 hydrophobic interactions. Figure 4. A. shows the binding interface of Amubarvimab and wild-type RBD, which is stabilized by a large number of hydrogen bonds(yellow). The Amubarvimab-N439K complex is stable for 12 Hydrogen bonds, 3 Electrostatic and 9 hydrophobic interactions, although the Hydrogen Bond number is not as abundant as that of the Wild-Amubarvimab complex, the number of interactions between Hydrophobic groups is higher than that of the Wild-Amubarvimab complex. Therefore, Amubarvimab also has a specific neutralizing ability when facing N439K mutation. Figure 5. B. shows the interaction between Amubarvimab and N439K mutation, with a reduced number of hydrogen bonds. The L452R-Amubarvimab complex is stabilized by 10 Hydrogen bonds, 2 Electrostatic and 8 hydrophobic interactions. Interestingly, Although the L452R-Amubarvimab complex is not dominant in the number of covalent bonds, the docking score of the L452R-Amubarvimab complex is higher than that of the N439K-Amubarvimab complex. It turns out that the weak hydrogen bond interaction between the L452R-Amubarvimab complex is less, and there is a solid Hydrophobic interaction (TYR489-LEU99). Instead, look at the two lowest-scoring compounds in the Amubarvimab group: E406W-Amubarvimab and Q498Y-Amubarvimab complexes. In the E406W-Amubarvimab complex, only 5 Hydrogen bonds and 6 Hydrophobic interaction, and there is no Electrostatic interaction at all. Moreover, the antigenic epitope of the E406W mutant RBD is minimal, and only 7 amino acid residues interact with Amubarvimab, and all of them are concentrated in a small area of the lower part of the RBD. The Q498Y-Amubarvimab complex, like the E406W-Amubarvimab complex, has very little non-covalent Bond interaction: only 7 Hydrogen bonds and 7 Hydrophobic interactions, and no Electrostatic interactions. Figure 5. D. and E. are the interaction modes of Amubarvimab with E406W and Q498Y, with few hydrogen bonds and no electrostatic interaction. The decrease in the number of interactions between Hydrogen Bonds and Hydrophobic and the lack of Electrostatic interactions cause the instability of these two complexes.

**Table 5.**
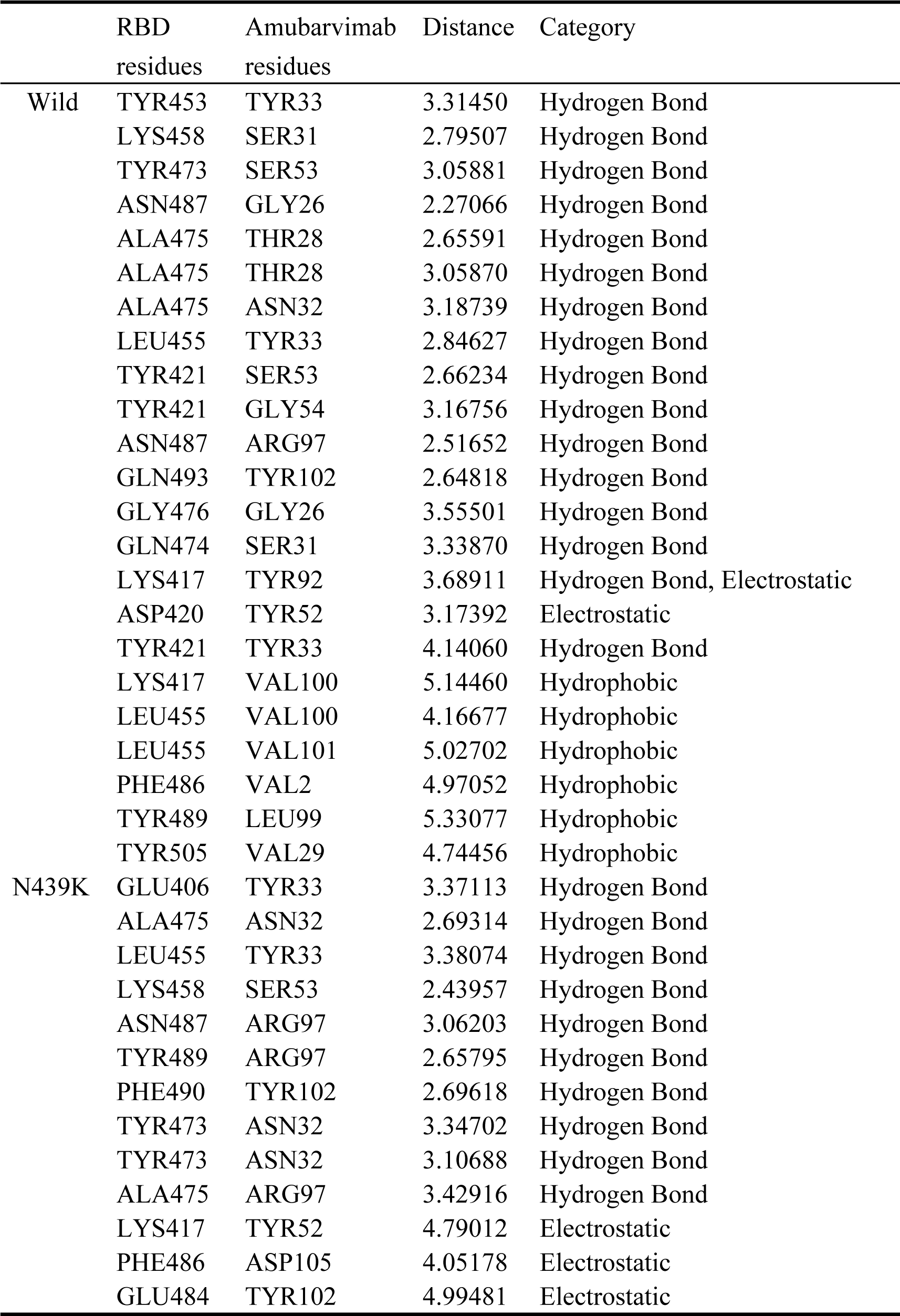
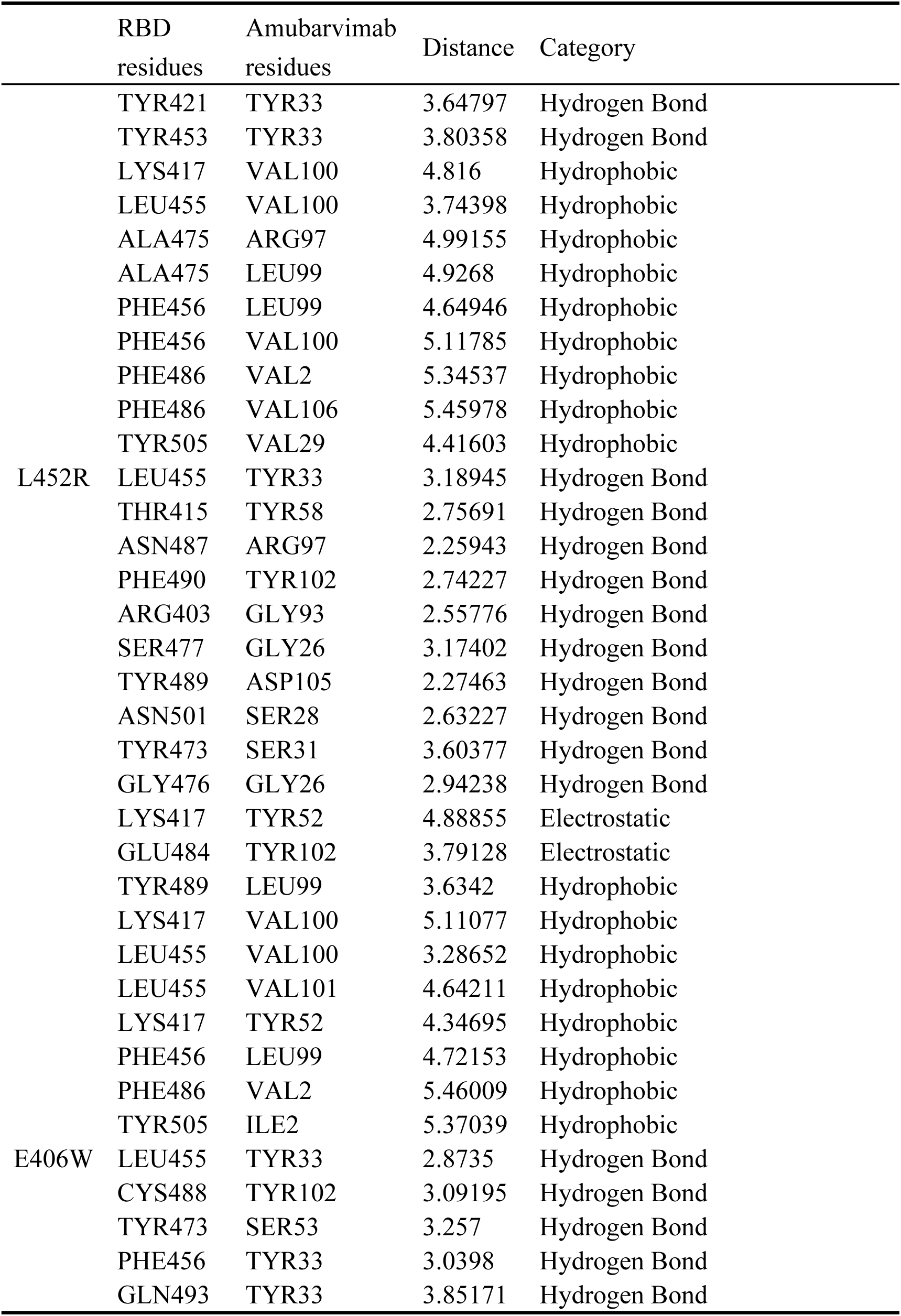
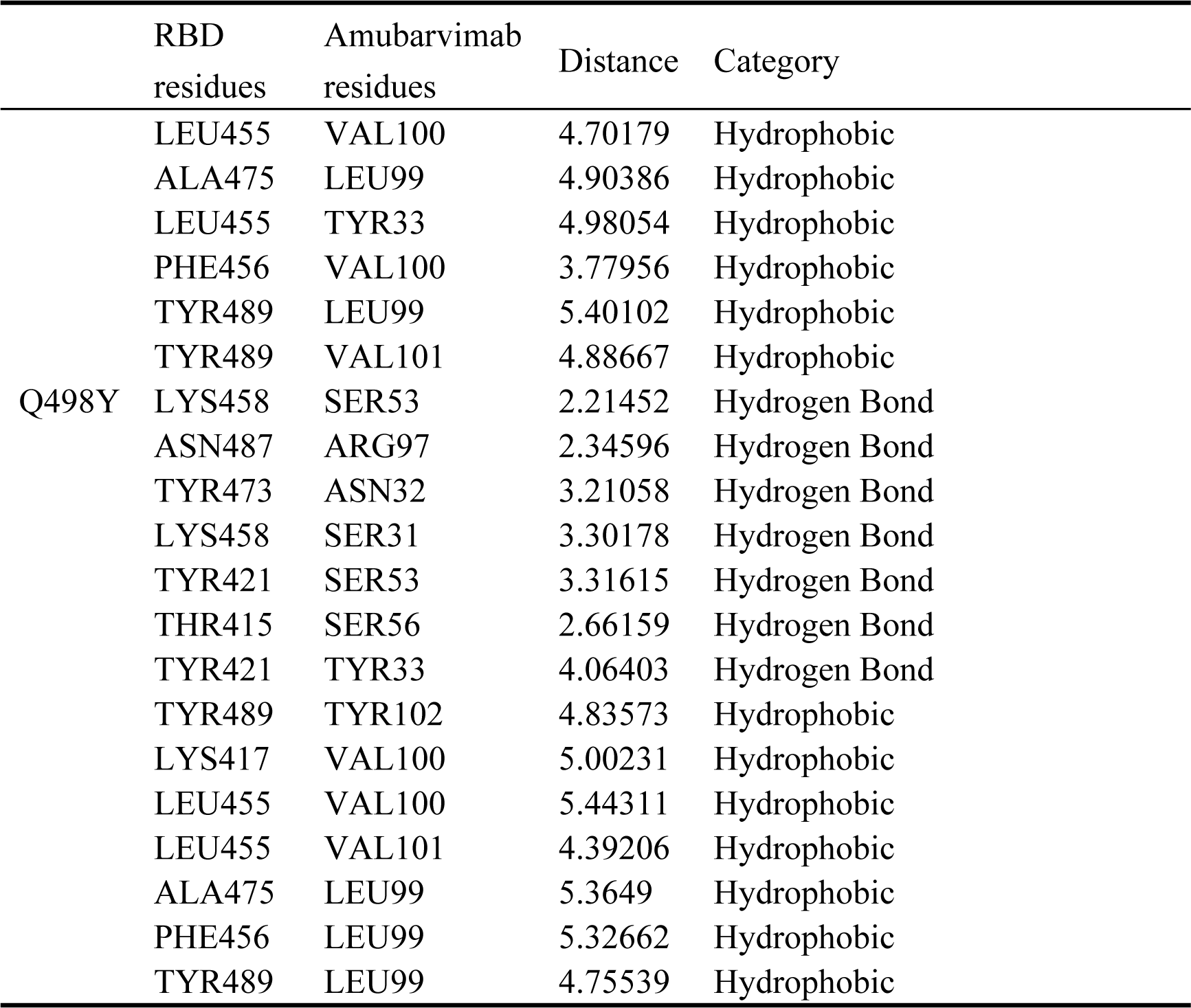
Biomolecular interaction analysis of Amubarvimab with wild-type and various mutant SARS-COV-2 RBDs.

**Figure. 5.**
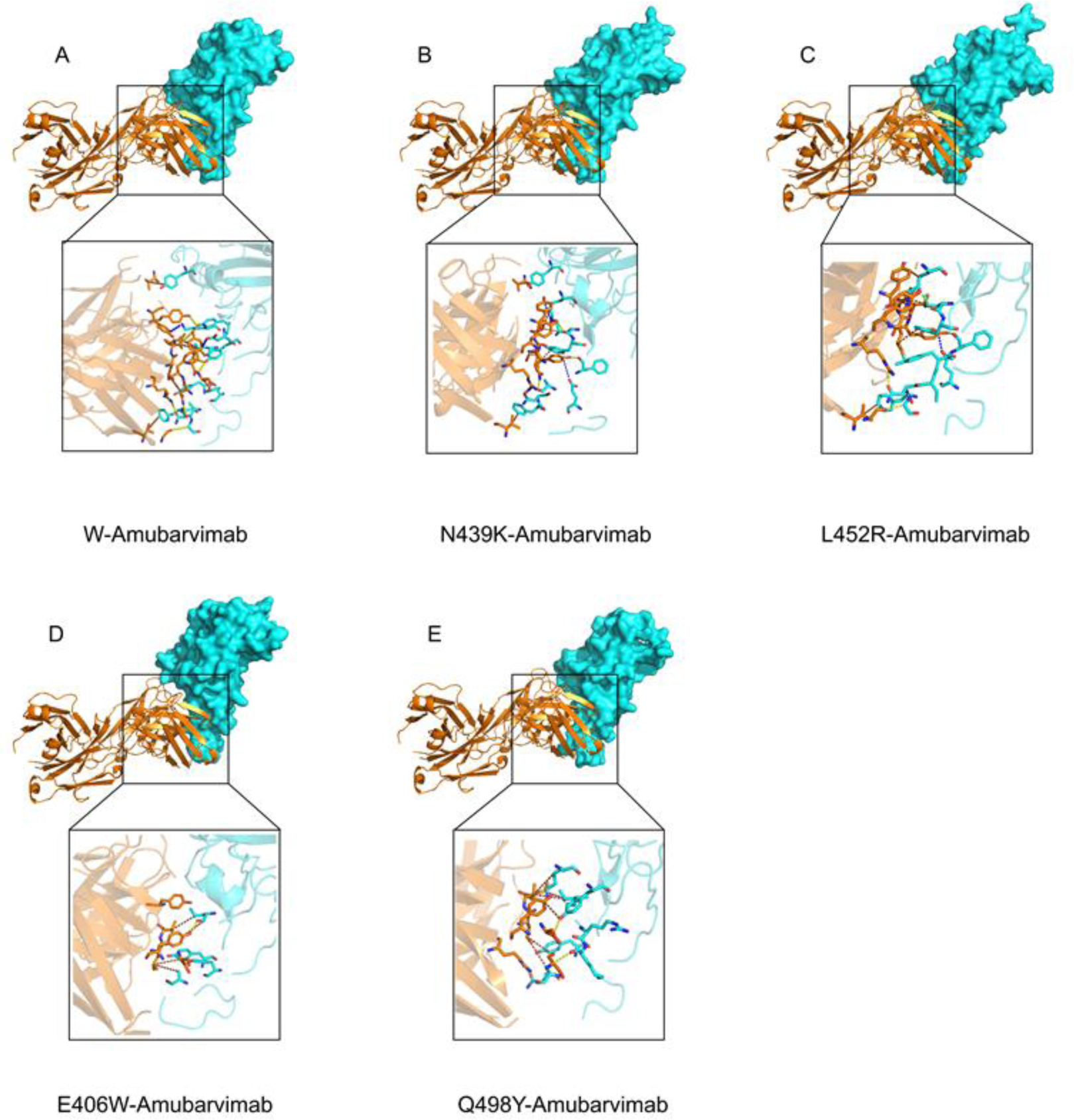
Schematic diagram of several SARS-COV-2 RBD combined with Amubarvimab. The blue surface model is RBD, and the yellow cartoon model is Amubarvimab. The neutral interface is enlarged to show the non-covalent interactions between the complexes, with hydrogen bonding interactions in yellow, electrostatic interactions in blue, and hydrophobic interactions in brown.

The Wild-Romlusevimab complex was used as the control to analyze the compound of the Romlusevimab group. As shown in Table 6, the Wild-Romlusevimab complex has 12 Hydrogen bonds, 4 Electrostatic and 6 Hydrophobic interactions, ensuring that Romlusevimab can effectively neutralize wild-type SARS-COV-2. The N439K-Romlusevimab complex showed a higher docking score, and it is worth noting that the N439K-Romlusevimab complex is inferior to the Wild-Romlusevimab complex in terms of the number of non-covalent bonds (9 Hydrogen bonds, 5 Electrostatic interactions and two hydrophobic interactions). But still showed high docking scores. After careful comparison, we found that the distance between the interacting amino acids in the N439K-Romlusevimab complex was generally close, while the distance between the interacting amino acids in Wild-Romlusevimab complex was generally far, which weakened the mutual right and left between amino acids. The N439K-Romlusevimab complex has more Electrostatic interactions, so the N439K-Romlusevimab complex shows a higher docking score. In the high-score complexes of Romlusevimab and various RBD interfaces, we found that many Electrostatic interactions between complexes maintain protein-protein binding, and it is speculated that Romlusevimab may effectively neutralize the RBD of SARS-COV-2 through its interactions. In Figure 6. B. and C. are the interaction modes of Romlusevimab and N439K, Y453F mutations. It can be seen that the electrostatic interaction increases, maintaining the stable conformation of the complexes. We subsequently verified this conjecture by comparing the low-scoring complexes of Romlusevimab and RBD. The interface scores of E406W-Romlusevimab and Q498Y-Romlusevimab are the lowest, only 11.34 and 11.7, and their Electrostatic interactions are only 2 and 3. When hydrogen bonds and hydrophobic interactions are not dominant, The Electrostatic interaction is reduced, which resulted in a weaker interaction between Romlusevimab and these two mutant RBD.

**Table 6.** Biomolecular interaction analysis of Romlusevimab with wild-type and various mutant SARS-COV-2 RBDs.

|  | RBD<br>residues | Romlusevimab<br>residues | Distance | Category |
| --- | --- | --- | --- | --- |
| Wild | ARG466 | ASP53 | 2.79217 | Hydrogen Bond, Electrostatic |
|  | ARG346 | GLU95 | 5.36007 | Electrostatic |
|  | ARG466 | ASP53 | 5.36179 | Electrostatic |
|  | THR345 | SER93 | 2.33177 | Hydrogen Bond |
|  | THR345 | SER93 | 3.33405 | Hydrogen Bond |
|  | ASN450 | ASN52 | 2.78397 | Hydrogen Bond |
|  | ASN360 | SER67 | 3.00667 | Hydrogen Bond |
|  | ASN448 | ASN53 | 2.33446 | Hydrogen Bond |
|  | ASN450 | ASN53 | 2.42235 | Hydrogen Bond |
|  | ALA352 | THR97 | 3.16766 | Hydrogen Bond |
|  | LYS356 | GLY29 | 3.60947 | Hydrogen Bond |
|  | ASN450 | THR31 | 2.53355 | Hydrogen Bond |
|  | ARG466 | TYR49 | 2.8886 | Hydrogen Bond |
|  | TYR449 | THR31 | 2.86886 | Hydrogen Bond |
|  | ARG466 | TYR50 | 3.62557 | Electrostatic |
|  | ALA348 | LEU99 | 4.12038 | Hydrophobic |
|  | ALA352 | ILE96 | 4.97051 | Hydrophobic |
|  | ALA352 | LEU99 | 3.92915 | Hydrophobic |
|  | TYR351 | ILE96 | 5.06814 | Hydrophobic |
|  | ARG355 | TYR50 | 4.88436 | Hydrophobic |
|  | ALA344 | TRP91 | 4.96916 | Hydrophobic |
|  | GLU340 | LYS31 | 2.47480 | Hydrogen Bond, Electrostatic |
| N439K | ARG346 | ASP96 | 5.15416 | Electrostatic |
|  | ARG357 | ASP51 | 3.53811 | Electrostatic |
|  | ARG466 | ASP53 | 4.85688 | Electrostatic |
|  | ASN360 | ASN27 | 3.24406 | Hydrogen Bond |
|  | ASN360 | SER67 | 2.25418 | Hydrogen Bond |
|  | SER359 | GLY68 | 3.21233 | Hydrogen Bond |
|  | ASN354 | THR98 | 2.95747 | Hydrogen Bond |
|  | SER359 | SER67 | 2.83135 | Hydrogen Bond |
|  | GLU340 | SER93 | 3.37163 | Hydrogen Bond |
|  | ARG357 | GLY29 | 3.15001 | Hydrogen Bond |
|  | SER359 | SER67 | 2.93476 | Hydrogen Bond |
|  | ARG466 | TYR50 | 3.47938 | Electrostatic |
|  | ILE468 | ILE96 | 4.76401 | Hydrophobic |
|  | ALA352 | LEU99 | 4.28457 | Hydrophobic |

**Table 6 continued Biomolecular interaction analysis of Romlusevimab with** **wild-type and various mutant SARS-COV-2 RBDs.**
|  | RBD<br>residues | Romlusevimab<br>residues | Distance | Category |
| --- | --- | --- | --- | --- |
| Y453F | GLU340 | LYS31 | 5.16098 | Electrostatic |
|  | ARG346 | ASP96 | 5.46111 | Electrostatic |
|  | ARG346 | ASP96 | 4.34506 | Electrostatic |
|  | LYS356 | ASP92 | 5.35201 | Electrostatic |
|  | ARG357 | ASP51 | 4.39448 | Electrostatic |
|  | ARG466 | ASP53 | 4.76040 | Electrostatic |
|  | SER359 | ASN27 | 2.35813 | Hydrogen Bond |
|  | ASN360 | ASN27 | 2.39168 | Hydrogen Bond |
|  | TYR351 | TYR32 | 2.98501 | Hydrogen Bond |
|  | THR345 | ILE94 | 3.38671 | Hydrogen Bond |
|  | ARG346 | THR98 | 2.97879 | Hydrogen Bond |
|  | ARG357 | LYS31 | 3.14365 | Hydrogen Bond |
|  | GLU340 | SER93 | 2.49729 | Hydrogen Bond |
|  | ARG346 | TRP91 | 4.70398 | Electrostatic |
|  | ARG466 | TYR50 | 4.06259 | Electrostatic |
|  | LYS356 | LYS31 | 5.06737 | Hydrophobic |
|  | ILE468 | ILE96 | 4.64854 | Hydrophobic |
|  | ALA344 | ILE94 | 4.56775 | Hydrophobic |
|  | ALA352 | ILE96 | 5.18928 | Hydrophobic |
| E406W | ARG357 | ASP51 | 4.93941 | Electrostatic |
|  | ARG355 | TYR50 | 2.65746 | Hydrogen Bond |
|  | GLU340 | SER93 | 2.68332 | Hydrogen Bond |
|  | THR470 | THR28 | 2.97138 | Hydrogen Bond |
|  | THR345 | ILE94 | 3.05402 | Hydrogen Bond |
|  | TYR351 | THR31 | 2.24283 | Hydrogen Bond |
|  | ASN360 | SER67 | 2.74591 | Hydrogen Bond |
|  | TYR449 | ASN56 | 3.35335 | Hydrogen Bond |
|  | ARG466 | THR97 | 2.66500 | Hydrogen Bond |
|  | ASN448 | THR54 | 3.25979 | Hydrogen Bond |
|  | ALA344 | ILE94 | 3.23126 | Hydrogen Bond |
|  | ARG346 | TRP50 | 4.30321 | Electrostatic |
|  | ARG357 | TYR50 | 5.45533 | Hydrophobic |
|  | ILE468 | TYR32 | 4.34134 | Hydrophobic |
|  | ARG346 | TRP50 | 5.10299 | Hydrophobic |
| Q498Y | ARG346 | GLU95 | 3.94798 | Hydrogen Bond, Electrostatic |
|  | LYS444 | ASP96 | 4.56852 | Electrostatic |

**Table 6 continued Biomolecular interaction analysis of Romlusevimab with** **wild-type and various mutant SARS-COV-2 RBDs.**
| RBD<br>residues | Romlusevimab<br>residues | Distance | Category |
| --- | --- | --- | --- |
| ARG466 | ASP53 | 5.45386 | Electrostatic |
| THR345 | SER32 | 3.22962 | Hydrogen Bond |
| ARG346 | SER32 | 3.27323 | Hydrogen Bond |
| ARG346 | TYR50 | 2.65790 | Hydrogen Bond |
| LEU492 | THR31 | 3.09052 | Hydrogen Bond |
| THR345 | SER30 | 3.19165 | Hydrogen Bond |
| THR345 | TRP91 | 3.22301 | Hydrogen Bond |
| ARG346 | HIS34 | 3.07599 | Hydrogen Bond |
| ASN450 | ASP96 | 2.42372 | Hydrogen Bond |
| ARG509 | SER93 | 3.20985 | Hydrogen Bond |
| LYS356 | SER52 | 3.72233 | Hydrogen Bond |
| SER494 | THR31 | 3.77848 | Hydrogen Bond |
| PHE490 | THR28 | 3.14492 | Hydrogen Bond |
| LEU441 | ILE94 | 5.16551 | Hydrophobic |
| ALA348 | TYR50 | 5.12551 | Hydrophobic |
| ARG346 | TRP91 | 4.50853 | Hydrophobic |
| LEU492 | TYR32 | 5.04817 | Hydrophobic |

**Figure. 6.**
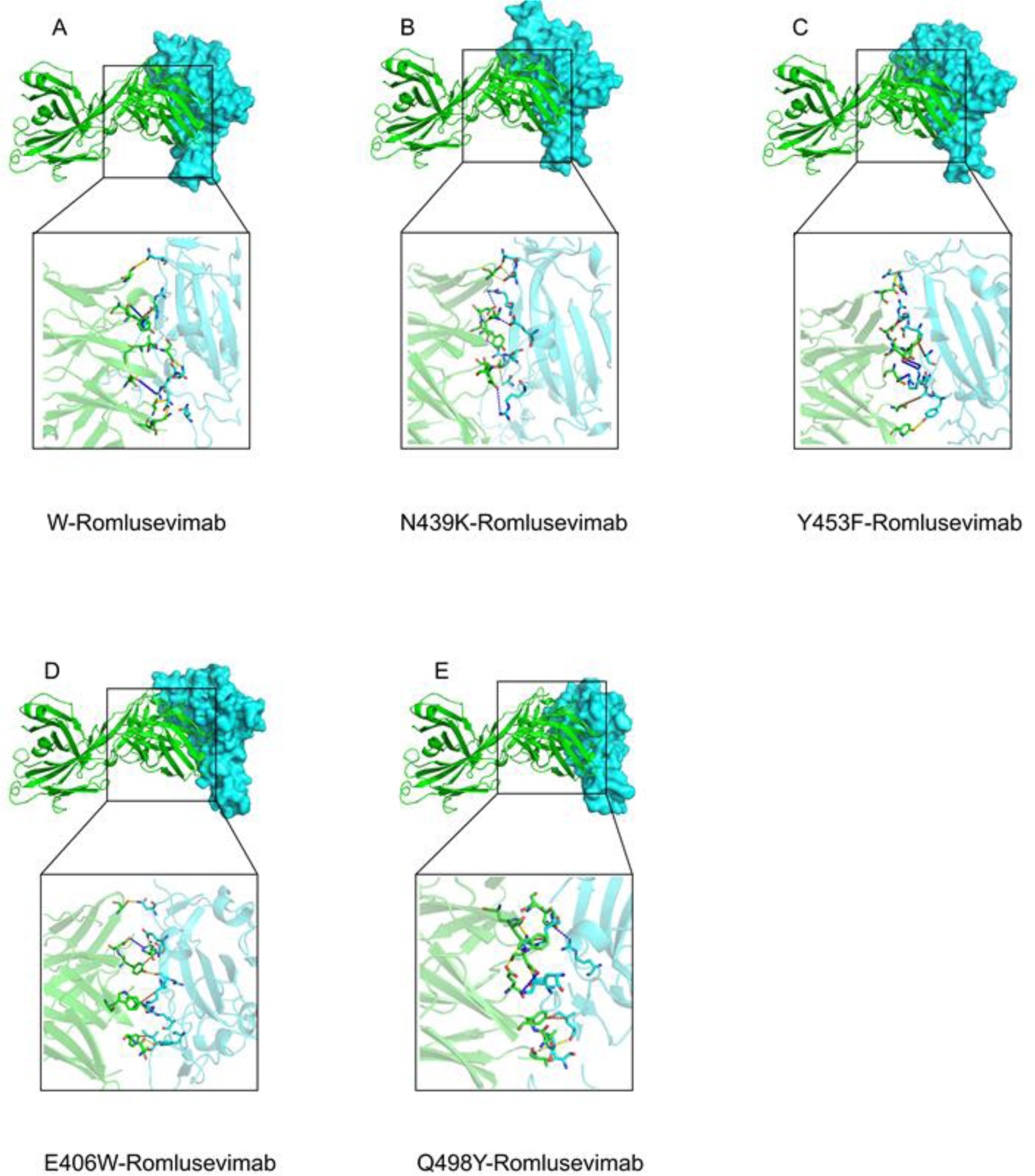
Schematic diagram of several SARS-COV-2 RBD combined with Romlusevimab. The blue surface model is RBD, and the green cartoon model is Romlusevimab. The neutral interface is enlarged to show the non-covalent interactions between the complexes, with hydrogen bonding interactions in yellow, electrostatic interactions in blue, and hydrophobic interactions in brown.

## Conclusion

In this study, we used protein molecular docking, Molecular dynamics simulation and interaction analysis techniques to explore the neutralization ability and neutralization mechanism of Amubarvimab and Romlusevimab against SARS-COV-2 and its variants. Our experiment proved that Amubarvimab has a high neutralizing effect on wild-type SARS-COV-2, moderate neutralization against the large majority of mutations from RBD, and low neutralization against the E406W and Q498Y mutations. Amubarvimab maintains the stability of the mAb-RBD complexes mainly through hydrogen bond interaction. When the number of hydrogen bonds decreases or the interaction distance increases, the stability of the complex will be significantly reduced. Romlusevimab also showed a certain neutralization effect on the SARS-COV-2 and its mutations, but its efficacy was not as good as that of Amubarvimab, especially in the face of E406W and Q498Y mutations, the docking score and the number of non-covalent bonds of complexes were much lower than other complexes. The electrostatic interaction plays an essential role in maintaining the stable conformation of the complexes, and there are a large number of electrostatic interactions in several strongly bound complexes. From the perspective of mixed antibodies, Amubarvimab and Romlusevimab bind to non-overlapping epitopes of RBD, which is non-competitive binding, and the combination of the two can prevent the immune escape of single mutations with a high probability, but in the face of E406W and Q498Y mutations, both of them show low binding ability. Therefore, antibodies mixed with Amubarvimab and Romlusevimab may show decreased neutralization efficacy in the face of E406W and Q498Y mutations.

## Acknowledgments

This work was financially supported by Natural Science Foundation of Shandong, China [Grant No. ZR2023MC059] and High-end Talent Introduction “Double Hundred Plan” Special Foundation of Yantai City [Grant No. 612211012002].

